# Structural Insights into the Activation and Inhibition of the ADAM17-iRhom2 Complex

**DOI:** 10.1101/2024.12.22.629961

**Authors:** Joseph J. Maciag, Conner Slone, Hala F. Alnajjar, Maria F. Rich, Bryce Guion, Carl P. Blobel, Tom C.M. Seegar

## Abstract

The endopeptidase activity of ADAM (a disintegrin and metalloprotease) -17, the primary processor of several EGFR ligands and tumor necrosis factor-alpha (TNF-α), is essential for proper embryonic development and immune regulation. Dysregulated ADAM17 activity is prevalent in a wide array of human diseases, including cancer, chronic inflammation, and SARS-CoV-2 viral progression. Initially translated as an inactive enzyme zymogen, ADAM17 maturation and enzymatic function are tightly regulated by its obligate binding partners, the inactive rhomboid proteins (iRhom) -1 and -2. Here, we present the cryo-EM structure of the ADAM17 zymogen bound to iRhom2. Our findings elucidate the interactions within the ADAM17-iRhom2 complex, the inhibitory mechanisms of the therapeutic MEDI3622 antibody and ADAM17 prodomain, and the previously unknown role of a membrane-proximal cytoplasmic re-entry loop of iRhom2 involved in the mechanism of activation. Importantly, we perform cellular assays to validate our structural findings and provide further insights into the functional implications of these interactions, paving the way for developing therapeutic strategies targeting this biomedically critical enzyme complex.

## INTRODUCTION

Ectodomain shedding is a biological process whereby proteolytic release of membrane-tethered proteins allows cells to communicate with one another and interpret their extracellular environment ^1^. The ADAM (a disintegrin and metalloprotease) family of membrane-anchored metalloproteases are type-1 transmembrane proteins with endopeptidase activity that participate in ectodomain shedding. ADAMs are widely expressed throughout the body with essential functions in fertilization, cell differentiation, angiogenesis, immunity and development of nervous and epithelial tissue ^2, 3^. Correspondingly, dysfunctional ADAM activity is thought to contribute to a variety of pathological conditions; such as auto-immune diseases, cancer and Alzheimer’s disease ^1, 4, 5, 6, 7^.

Among the ADAM family, ADAM17 stands out for its essential roles in EGFR-signaling during embryonic development, adult homeostasis and disease pathogenesis ^2, 8, 9, 10^. ADAM17 was originally identified as the enzyme responsible for the release of soluble tumor necrosis factor-α (TNFα), a potent inflammatory cytokine, from cells ^11, 12^. Genetic ablation of ADAM17 in mice results in perinatal lethality, with mutant mice displaying open eyes at birth and defects in heart valve and growth plate formation, which has been attributed to a loss in epidermal growth factor receptor (EGFR) signaling ^8, 9, 13, 14, 15^. Patients lacking ADAM17 have severe skin and intestinal barrier defects ^16, 17^, likely caused by defects in the activation of EGFR-ligands ^18, 19^. Accordingly, ADAM17 was identified as the enzyme responsible for the maturation of several pro-EGF ligands into soluble signaling molecules ^8, 9, 10, 20, 21^. In addition to its function as the TNFα convertase, ADAM17 also cleaves a myriad of other membrane tethered proteins to shed them from the cell membranes ^3^, including the interleukin 6 receptor (IL-6R) ^5, 8, 13, 22, 23^, a key player in autoimmune disease.

During biosynthesis, ADAM17 is translocated into the endoplasmic reticulum (ER) as a zymogen, with a pro-domain tightly bound to its catalytic domain to maintain enzyme latency and prevent premature substrate processing ^24^. Maturation of ADAM17 from its zymogen form, a prerequisite for peptidase activity, involves the proteolytic processing of the prodomain by pro-protein convertases in the trans-Golgi network ^24, 25^. The mature ADAM17 contains an extracellular metalloprotease (M), ancillary disintegrin (D) and cysteine-rich (C) domain, a single transmembrane (TM) α-helix and a cytoplasmic domain (Figure 1) ^11, 12^.

**Figure 1:**
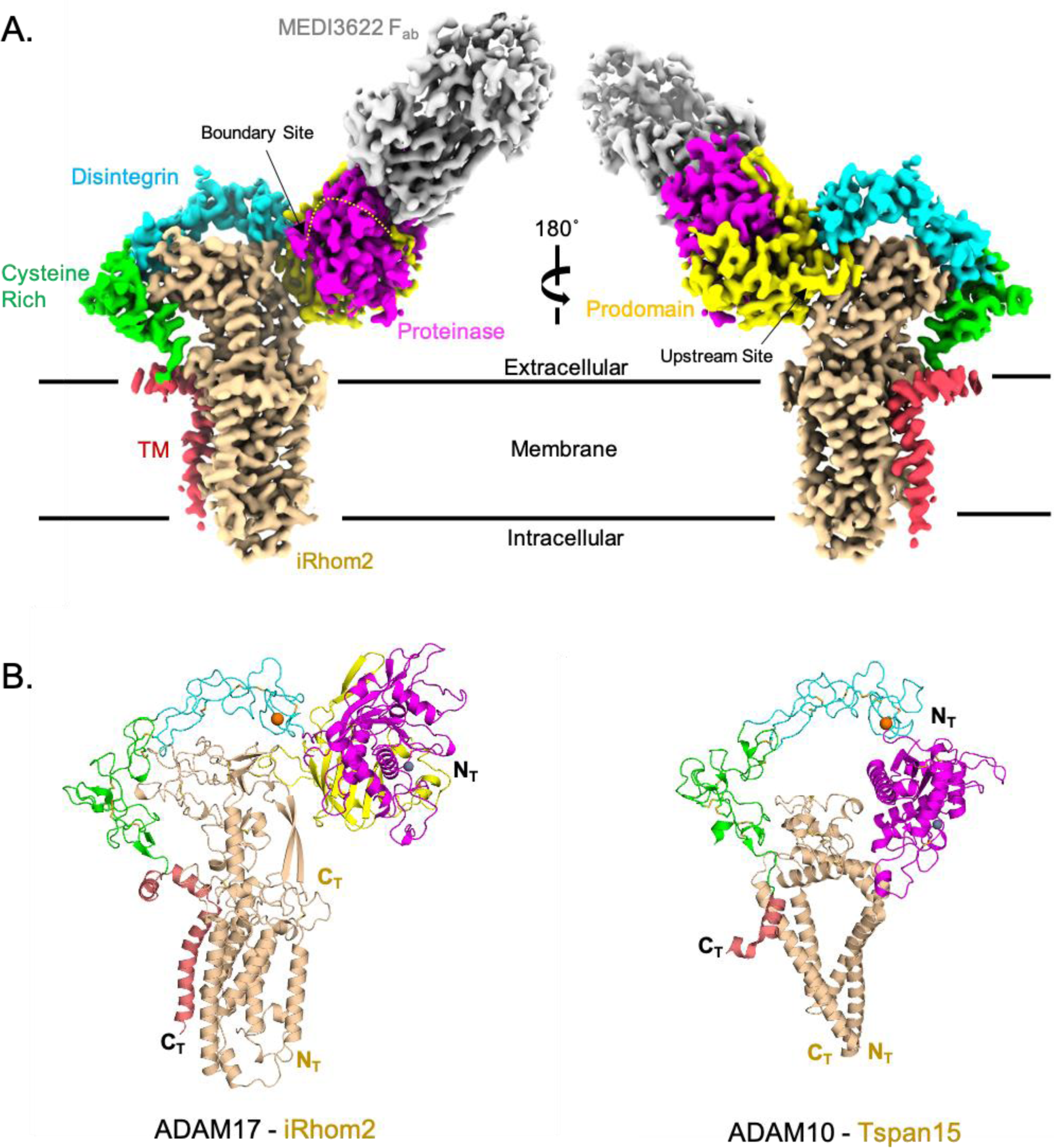
Overall Structure of the iRhom2-ADAM17 Zymogen Complex. **(A)** Cryo-EM density map of the zymogen ADAM17-iRhom2-MEDI3622 F_ab_ complex embedded within the cell membrane, represented by parallel black lines. ADAM17 is colored according to its structural domains: Prodomain (yellow), Protease (magenta), Disintegrin (cyan), Cysteine-Rich (green), and TM domains (red). iRhom2 is shown in tan, and MEDI3622 F_ab_ is depicted in gray. Proprotein convertase cleavage sites within the prodomain are indicated by arrows, denoting the Upstream (US) and Boundary (BS) cleavage sites. **(B)** Cartoon representation comparing the zymogen ADAM17-iRhom2 complex (Left) and the ADAM10-TSPAN15 complex (Right, RCSB PDB: 8ESV). ADAM17 and ADAM10 structures were superimposed based on their D and C domains and then separated for visualization. Structural domains are color-coded as in (A), with additional representation of calcium ions (orange) and catalytic zinc ions (dark gray) as spheres.

ADAM17 activity can be rapidly stimulated, with substrate shedding induced within minutes of activation ^20, 26, 27, 28^. The short temporal activation and inability to increase enzyme activity by systemic overexpression in mice ^29^ highlight the posttranslational nature of the mechanism governing ADAM17 activation. While the ADAM17 cytoplasmic domain is required for normal mouse development ^30^, paradoxically, stimulated ADAM17-dependent shedding occurs independent of its cytoplasmic domain ^26, 31^ but requires the presence of its transmembrane domain ^28^. These observations underscored the likely importance of one or more other integral membrane proteins in regulating ADAM17 activity. To this end, the discovery of the seven-transmembrane inactive rhomboid proteins (iRhom) -1 and -2 as essential regulators of ADAM17 revealed the identity of its crucial regulatory binding partners ^32, 33, 34, 35^.

iRhoms are part of the rhomboid protease superfamily of proteins that have evolved to include a large extracellular domain and have lost the amino acids required to function as an integral membrane protease, making them functionally distinct from other rhomboid protease family members ^36, 37, 38^. Mice lacking iRhom2 fail to generate mature and functional ADAM17 in hematopoietic lineage cells and therefore release little, if any, soluble TNFα from stimulated immune cells in proinflammatory conditions ^32, 33^. Yet, unlike *Adam17^−/−^* mice, which die shortly after birth, mice lacking iRhom2 have no spontaneous pathological phenotypes ^5, 32, 33^. However, double knockout mice lacking *iRhom1* and *iRhom2* die shortly after birth with open eyes, heart valve and growth plate defects, thus closely resembling mice lacking *Adam17* ^35^. No mature, functional ADAM17 can be detected in *iRhom1/2^−/−^* cells and tissues, including a second *iRhom1/2-*deficient mouse strain that dies during embryogenesis ^34^. These findings highlight the essential role iRhoms in the maturation and function of ADAM17. Moreover, these studies suggest that either iRhom can bind ADAM17 zymogen in the ER to facilitate its exit from the ER ^34, 35^, which allows the iRhom/ADAM17 complex to undergo maturation in the trans-Golgi network. iRhoms are also thought to accompany ADAM17 *en route* to the cell surface, where they regulate its extracellular peptidase activity ^39, 40, 41, 42, 43, 44^. A point mutation in the TM1 of iRhom2 identified in mice, termed *sinecure*, strongly reduces the maturation and function of ADAM17, which established the importance of the iRhom2 TM1 as an essential component of the regulation of ADAM17 ^45, 46^. Replacing the TM of ADAM17 with that of betacellulin gives rise to a constitutively active protease that can no longer be rapidly stimulated and does not require the iRhoms for its catalytic activity ^28, 47^. Moreover, cells lacking ADAM17 have little, if any iRhom2 ^48^, whereas immune cells lacking iRhom2 have no mature ADAM17 ^32, 33^, establishing a reciprocal dependence of both binding partners for their maturation and function. Thus, iRhoms are considered vital regulators of ADAM17 activity.

Here, we report a high-resolution cryo-EM structure of the ADAM17 zymogen bound to iRhom2 and the MEDI3622 therapeutic antibody and the results of structurally informed cell-based assays to identify key structural features of the complex essential for ADAM17 activity. Critically, beyond reinforcing recent structural work ^49^, we identify a membrane-proximal cytoplasmic region in iRhom2 that integrates into its transmembrane α-helix bundle and is crucial for ADAM17 sheddase function. Additionally, we provide atomic insights into the inhibitory mechanisms of the MEDI3622 therapeutic antibody and the ADAM17 prodomain in blocking ADAM17 catalytic activity. These discoveries deepen our understanding of ADAM17 regulation and function, and will inform the development of novel therapeutic strategies targeting this key enzyme complex.

## RESULTS

### Structure of zymogen ADAM17 – iRhom2 Complex

To establish optimal expression and isolation procedures for structural studies, we first created a series of iRhom2 cytoplasmic deletion constructs, replacing portions of this domain with the fluorescent reporter protein mVenus. The iRhom2 cytoplasmic domain is not predicted to bind the ADAM17 cytoplasmic domain, a region dispensable for stimulated endopeptidase activity ^26, 28, 50^. The chimeric mVenus-iRhom2 proteins, along with ADAM17, were expressed in Expi293 cells, detergent solubilized and screened using fluorescent-based size exclusion chromatography (FSEC) to identify optimal conditions to isolate the ADAM17-iRhom2 complex (Supplementary Figure 1A and 1B). Interestingly, iRhom2 mutants with truncated forms of the cytoplasmic domain were less susceptible to sample degradation than full length iRhom2 and still retained the ability to bind ADAM17 (Supplementary Figure 1A).

To determine the structure of the complex, the mVenus-Δ363-iRhom2 cytoplasmic deletion construct and ADAM17 with the inactivating protease mutation E406A were co-expressed in Expi293 cells. The complex was solubilized in detergent buffer and purified using an αGFP-nanobody affinity column. The purified zymogen ADAM17-iRhom2 complex was then bound by a F_ab_ fragment, corresponding to a function blocking α-ADAM17 antibody (MEDI3622), ^51^ to learn more about its mechanism of inhibition and to serve as a fiducial marker for cryo-EM imaging and structure determination (Supplementary 1C). The structure of the zymogen ADAM17-MEDI3622 F_ab_ complex bound to iRhom2 was determined using single particle analysis to a global resolution of 3.53 Å, with a local resolution minimum of 1.93 Å (Figure 1 and Supplementary Figure 1D). The final density map displayed clear structural features for zymogen ADAM17 and iRhom2 embedded in a glycol-diosgenin (GDN) micelle, with the MEDI3622 F_ab_ bound to the ADAM17 extracellular domain (ECD). However, no density was observed for the cytoplasmic regions of ADAM17 or the iRhom2-mVenus fusion, likely due to native disorder and high flexibility in the cytoplasmic domains. To build the initial structure, AlphaFold predictive models for the MEDI3622 F_ab_, the Δ363-iRhom2 cytoplasmic deletion mutant and individual domains of ADAM17 were docked into the final density map, manually rebuilt, and refined (Table 1). Manual inspection of the final protein model in the density map showed good correlation between the Cα-backbone and placement of large branched amino acid side chains into density extending from the secondary structural elements.

**Table 1:**
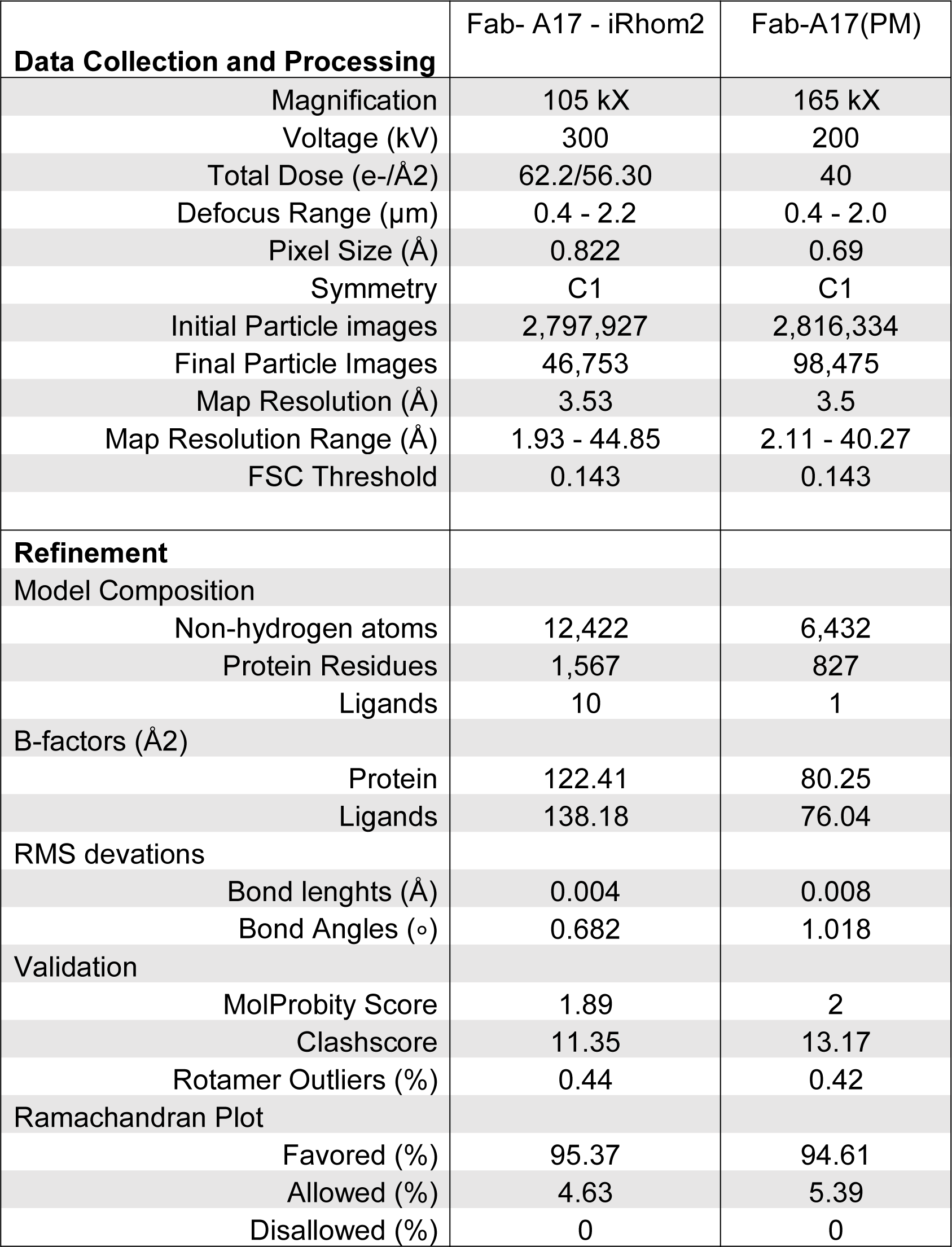
Cryo-EM Data Collection and Refinement Statistics.

The overall architecture of the zymogen ADAM17-iRhom2 complex reveals an extended conformation of the ADAM17 ECD, positioned proximal to the cell membrane making extensive contact with the iRhom2 ECD, known as the iRhom homology domain (IHD) (Figure 1). Both proteins are further stabilized and anchored in the membrane by their TM α-helices. The inhibitory MEDI3622 F_ab_ projects from the ECDs, bound to the ADAM17 metalloprotease (M) domain. Remarkably, the overall structure of zADAM17 bound to iRhom2 resembles the conformation adopted by ADAM17’s closest homolog, ADAM10, bound to the multi-pass transmembrane tetraspanin 15 (Tspan15) protein (RCSB PDB: 8ESV) ^52^. Superimposition of the ADAM17 ancillary D+C domains onto ADAM10 (RMSD = 2.65 Å) positions the transmembrane binding partners, Tspan15 and iRhom2, as a molecular bridge between the ADAM C and M domains (Figure 1B). Unlike the ADAM10-Tspan15 structure, the ADAM17-iRhom2 complex shares extensive contact within the transmembrane regions.

### Inhibition of ADAM17

The ADAM17 M domain is divided into N-terminal (N_T_) and C-terminal (C_T_) lobes (Figure 2A), separated by a central catalytic cleft that use enzyme pockets, denoted as S, to position an extended protein substrate for proteolysis. The S positions are numbered relative to the scissile bond location in the protein substrate, with positions toward the N-terminal given unprimed notation (e.g., S1, S2), and those toward the C-terminal given primed notation (e.g., S1’, S2’) (Figure 2A). The active site of ADAM17 harbors a zinc ion coordinated by three histidine residues, along with a glutamic acid residue and a water molecule, which together facilitate the hydrolysis of a protein substrate between the S1 and S1’ recognition pockets ^53^. In our zymogen ADAM17-iRhom2 structure, the prodomain is observed bound to the ADAM17 M domain, wedged between this domain and the iRhom2 IHD (Figure1 and 2). Unlike other protease prodomains, which are typically short polypeptides, the ADAM17 prodomain is considerably larger, approximately 25 kDa, with no homologous structure available in the RCSB database. The ADAM17 prodomain folds into two stacked β-sheets, forming a β-barrel motif (β1-3 and β5-10) that binds to the M domain opposite the active site cleft, burying a substantial contact interface of 5380 Å² (Figure 2). Additional non-catalytic cleft contacts extend the central β-sheet in the M domain by two β-strands (β3 and β4). The C-terminal end of the prodomain, referred to as the “cysteine-switch,” contains a single cysteine residue (C184) whose thiol side chain coordinates the active site zinc ion. Though the C184 residue itself is dispensable for inhibition in cellular assays and in studies with using recombinant ADAM prodomain ^54, 55^, the region preceding C184 also occupies the catalytic cleft and is critical for inhibiting ADAM17 activity (Figure 2). To summarize, prodomain interactions inhibit ADAM17 by displacing the nucleophilic water molecule and sterically blocking substrate access.

**Figure 2:**
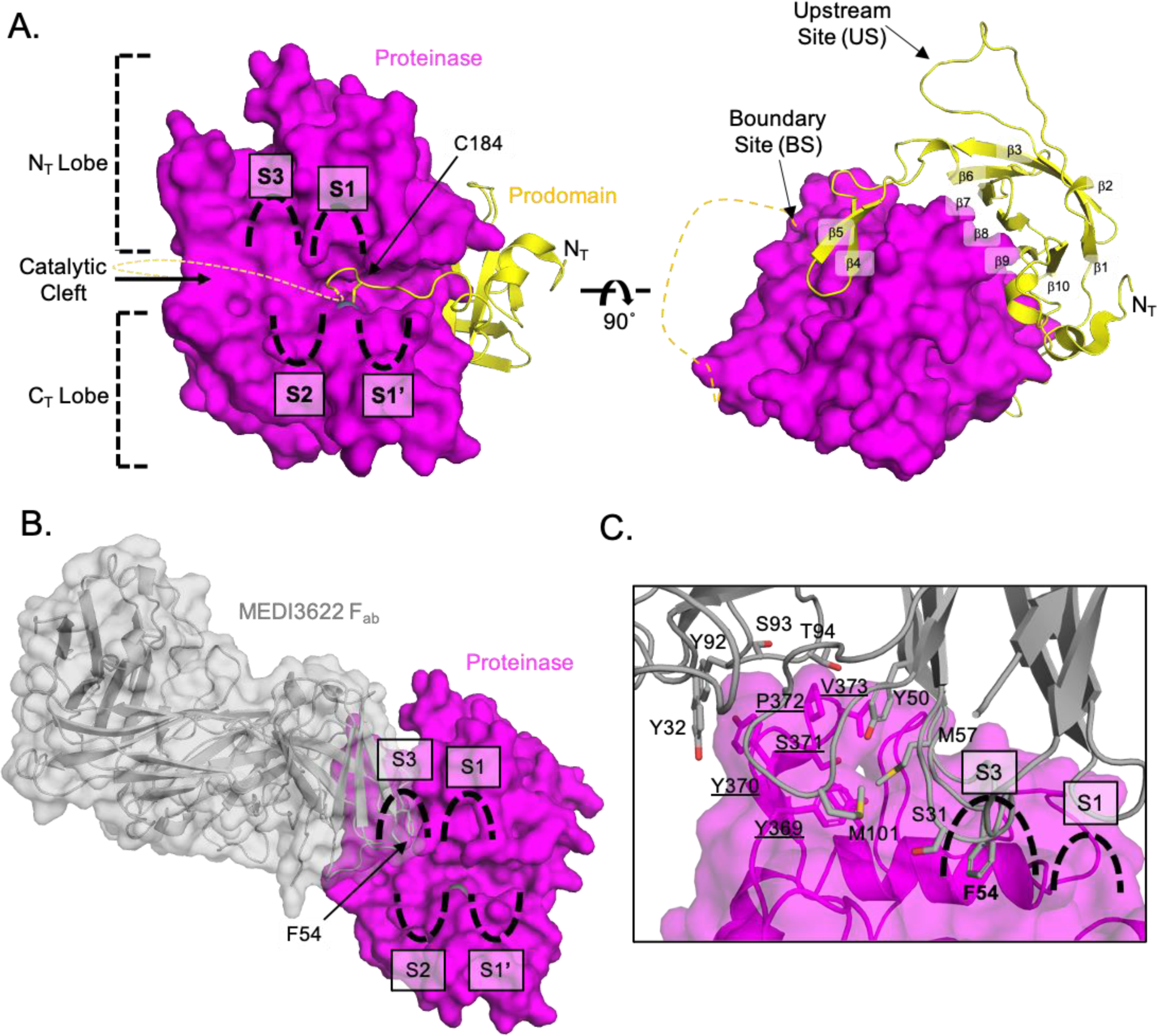
Structural Analysis for the Inhibition of ADAM17. **(A)** Cartoon representation of the prodomain and M domain of ADAM17, with the M domain surface shown in transparency. The M domain is divided into two lobes, N_T_ and C_T_ (dashed brackets), based on their positions relative to the central catalytic cleft containing the substrate-selective S3, S2, S1, and S1’ pockets (outlined). Structural β-strands (β1-10) of the prodomain and the cysteine switch residue, C184 (side chain depicted as sticks with RGB coloring), are annotated. The C_T_ region of the prodomain, including the BS cleavage sites, is represented as a dashed line. **(B)** Cartoon and surface representation of the MEDI3622 F_ab_ (gray) bound to the ADAM17 M domain, highlighting F52 of the MEDI3622 F_ab_, which occupies the S3 substrate-selective binding pocket. **(C)** Zoomed-in view of the amino acids within the contact interface between the MEDI3622 F_ab_ and the ADAM17 M domain (underlined).

Additionally, our structure elucidates the precise molecular mechanism of inhibition by the therapeutic antibody MEDI3622, demonstrating how its binding to the ADAM17 M domain obstructs substrate access and prevents proteolytic activity (Figure 2B). The complementarity-determining regions (CDRs) of the MEDI3622 F_ab_ binds to a loop, distal from the active site, in the ADAM17 M domain, specifically at amino acids Y369/Y370/S371, confirming the binding epitope previously identified by mutagenesis (Figure 2B-C)^56^. Substrate selectivity studies for ADAM10 and ADAM17 indicate a preference for amino acids up to four residues from the scissile bond, particularly favoring branched amino acids in the P3 position ^57, 58^. In support of this, the X-ray structure of ADAM10 shows a peptide product occupying the S1-S3 substrate-selective pockets ^59^. Notably, the placement of the F54 side chain in the CDR2 of the MEDI3622 F_ab_ heavy chain into the ADAM17 hydrophobic S3 substrate-selective pocket sterically obstructs an incoming protein substrate, which is crucial for ADAM17 activity. Moreover, the binding of MEDI3622 likely accounts for the lack of density for the prodomain region between the cysteine-switch and the Furin processing site in our structures. This region likely binds to the S3 pocket, as these amino acids have been implicated in the inhibitory properties of the ADAM17 prodomain ^54^. Altogether, the zymogen ADAM17-MEDI3622 cryo-EM structures provide atomic-level details of how MEDI3622 inhibits ADAM17 activity, effectively preventing access of the substrate to the catalytic cleft.

### Overall Structure of the iRhom2

iRhom2 has evolved from the rhomboid superfamily into a pseudoenzyme that lacks enzyme activity and features a large extracellular IHD. The IHD is a section of 243 amino acids nestled between TM1 and TM2, anchored into the cell membrane by an amphipathic α-helix preceding TM2 (Figure 3). This domain is stabilized by a network of 8 disulfide bonds, extending 45 Å from the cell surface, and capped by an unstructured region responsible for tethering the ADAM17 ECD (Figure 3A). The iRhom2 TM1-6 α-helices fold into a rhomboid-like domain with TM7 located to the periphery of the rhomboid-like α-helical bundle (Figure 3). The prototypical rhomboid protease, GlpG (RCSB PDB: 2IC8), superimposes well (RMSD = 2.36 Å) onto the core iRhom2 TM1-4 and TM6 (Supplementary Figure 3A). Interestingly, the iRhom2 TM5 is displaced from the core rhomboid fold by 12 Å to accommodate the insertion of a conserved iRhom2 cytoplasmic amphipathic α-helix (Supplementary Figure 3B), dubbed the “re-entry” loop, into the base of the rhomboid-like fold (Figure 3D).

**Figure 3:**
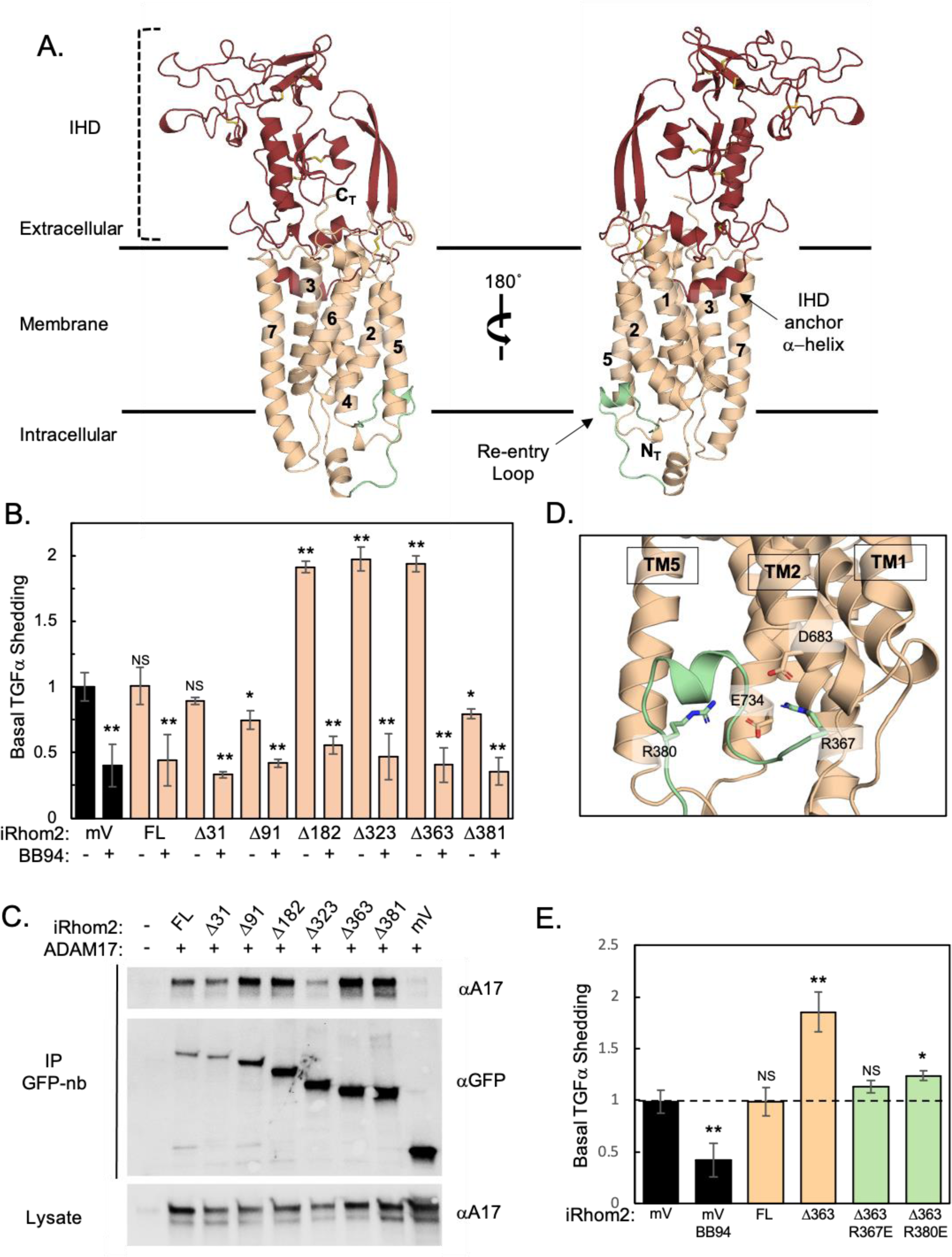
Analysis of the iRhom2 Cytoplasmic and Re-entry Loop in ADAM17 Activity. **(A)** Cartoon representation of iRhom2 within the cell membrane. The iRhom homology domain (IHD) is highlighted in crimson and indicated with a dashed bracket. The transmembrane (TM) α-helices are numbered sequentially (1–7), and the re-entry loop, shown in green, is marked with an arrow positioned between TM2 and TM5. **(B)** Basal shedding assay of TGFα mediated by ADAM17. U2OS cells were co-transfected with AP-TGFα and either mVenus (mV), full-length iRhom2 (FL), or iRhom2 constructs with cytoplasmic truncations. Normalized AP-TGFα shedding (N ≥ 3) was quantified with and without the pan-metalloprotease inhibitor BB94. All iRhom2 transfections were normalized to the mVenus control in the absence of BB94. **(C)** Immunoprecipitation (IP) analysis of mVenus-iRhom2 constructs with ADAM17. Expi293F cells were co-transfected with catalytically inactive ADAM17 E406A and either mVenus (mV), full-length iRhom2 (FL), or iRhom2 constructs with cytoplasmic truncations. Lysates immunoprecipitated using GFP nanobody (GFP-nb) resin were analyzed by western blot for ADAM17 (top panel) and iRhom2-mVenus (middle panel). Total ADAM17 levels in cell lysates were confirmed by western blot (bottom panel). **(D)** Close-up view of the re-entry loop with the residues R367, R380, D683, and E734 displayed as sticks and color-coded in RGB. **(E)** Basal shedding assay of TGFα mediated by ADAM17 with re-entry loop mutations. U2OS cells were transfected with AP-TGFα and either mVenus (mV), full-length iRhom2 (FL), Δ363-iRhom2, Δ363-iRhom2 R367E, or Δ363-iRhom2 R380E. AP-TGFα shedding (N ≥ 3) was normalized to the mVenus control. Error bars represent mean ± SD of independent experiments. Statistical significance was determined using an unpaired, two-tailed t-test between the mV sample and each iRhom2 sample. *p < 0.05; **p < 0.005; NS, not significant.

The cytoplasmic domain of iRhom2 is crucial for regulating the stimulated shedding of ADAM17 substrates from the cell surface. To evaluate the role of the re-entry loop in ADAM17 activity, we tested cytoplasmic deletion constructs of iRhom2 in a well-established cell-based assay measuring ADAM17 processing of the chimeric alkaline phosphatase-transforming growth factor α (AP-TGFα) reporter ^20, 60^ (Figure 3B-E). In these assays, AP-TGFα shedding was assessed in the presence and absence of the metalloprotease inhibitor BB94 to confirm the contribution of ADAM17 activity to basal shedding. Deleting the cytoplasmic region of iRhom2, specifically up to amino acid residue M183 (Δ182), led to an increase in basal shedding of AP-TGFα compared to the overexpression of full-length iRhom2 (Figure 3B), which is also seen with shorter deletion mutants (Δ323, Δ363). In contrast, deletion of the cytoplasmic region, including the re-entry loop (Δ381), nearly abolished basal shedding of AP-TGFα (Figure 3B), suggesting that the highly conserved re-entry loop sequences between R364 and Y382 are crucial for the regulation of ADAM17 (Figure 3D). Within the re-entry loop, R380 and R367, which stabilize the re-entry loop through salt bridge bonds with E733 and D682, are highly conserved and likely functionally important. To test this idea, the R367E and R380E mutations, which reverse the charge potential, attenuates the increased basal shedding activity observed with the Δ363-iRhom2 construct (Figure 3E). Importantly, mutations within the re-entry loop or its complete removal do not affect ADAM17 binding in co-immunoprecipitation experiments (Figure 3C), underscoring the re-entry loop of iRhom2 as a key cytoplasmic regulatory element that controls extracellular ADAM17-dependent substrate processing.

### Structure-function Studies of the ADAM17-iRhom2 Contact Interface

The ADAM17-iRhom2 complex is stabilized by non-covalent interactions that can be divided into two distinct regions: one involving the ECDs and the other between the transmembrane α-helices within the cell membrane (Figure 4A). Analysis of iRhom2’s surface charge potential revealed that the ECD is dominated by ionic interactions, notably through a negatively charged region within the IHD, proximal to a cluster of oppositely charged residues (R625, K626, and K628) in the ADAM17 C domain (Figure 4A Site2 and Supplementary Figure 4C). These amino acids in ADAM17 are functionally significant, as they were previously implicated in controlling ADAM17 enzyme activity required for normal embryonic development in mice ^61^.

**Figure 4:**
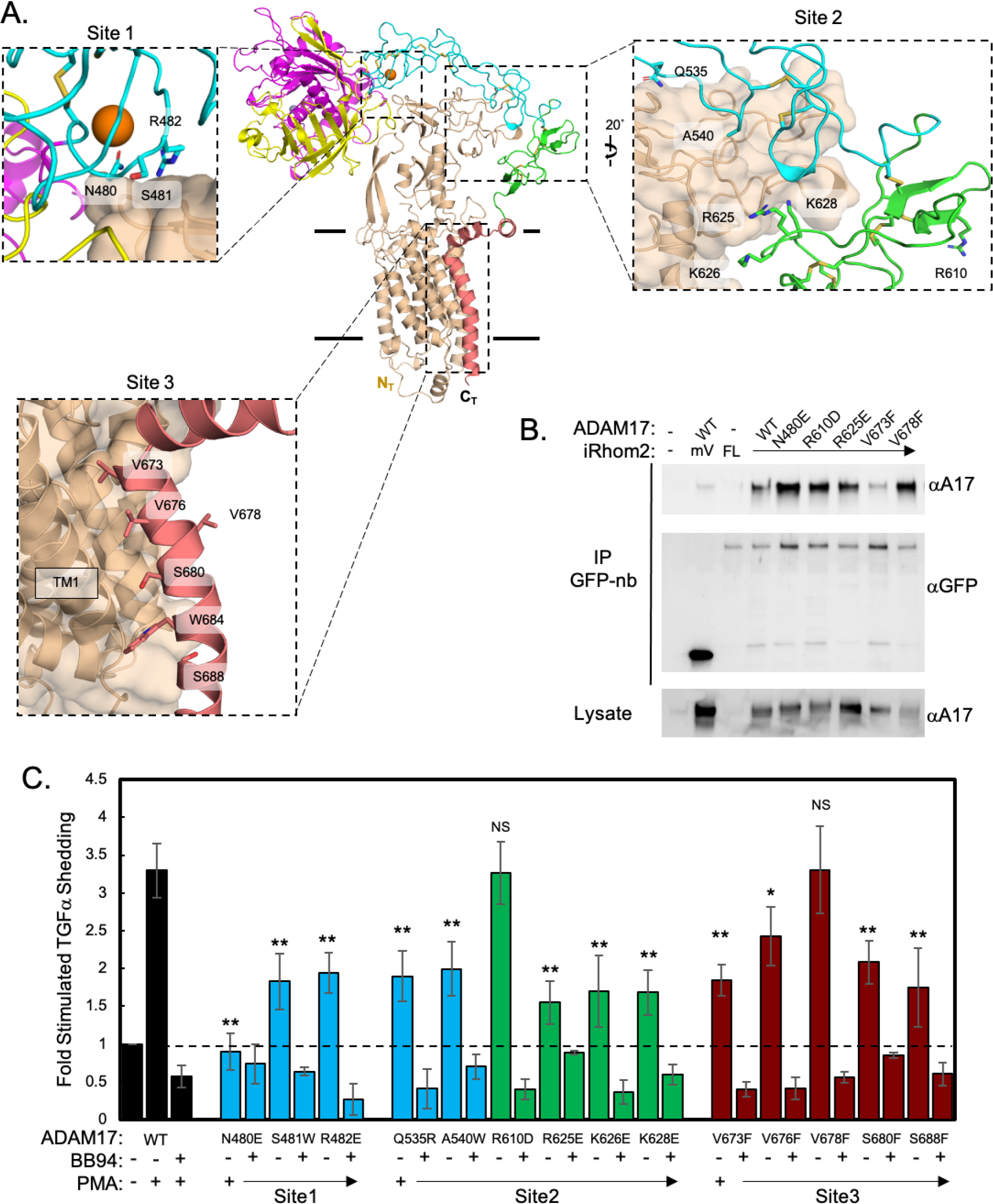
Analysis of the iRhom2-ADAM17 Contact Interface in ADAM17 Activity. **(A)** Cartoon representation of the ADAM17-iRhom2 complex showing three contact interfaces, designated as Sites 1–3. Zoomed views (inset boxes) highlight the amino acid residues at each interface, with side chains displayed as sticks. These residues were assayed for their effects on ADAM17 activity. **(B)** Immunoprecipitation (IP) analysis of mVenus-iRhom2 interaction with ADAM17 mutations. Expi293F cells were co-transfected with either mVenus, full-length mVenus-iRhom2, or ADAM17 constructs (WT or Site 1–3 mutants). Lysates were immunoprecipitated using GFP nanobody (GFP-nb) resin and analyzed by western blotting for ADAM17 (top panel) and mVenus-iRhom2 (middle panel). Total ADAM17 levels in lysates were confirmed by western blot (bottom panel). **(C)** Stimulated shedding assay of TGFα mediated by ADAM17. *Adam17^−/−^* cells were co-transfected with AP-TGFα and either WT ADAM17 or ADAM17 mutants. PMA-stimulated and BB94-treated samples were normalized to the unstimulated sample for each ADAM17 variant. Data are color-coded as follows: WT ADAM17 (black), D domain mutations (cyan), C domain mutations (green), and TM domain mutations (crimson). Error bars represent mean ± SD from *N* ≥ 3 independent experiments. Statistical significance, comparing PMA-stimulated samples, was determined using an unpaired, two-tailed t-test to compare individual mutant samples to the WT control under stimulated conditions. *p < 0.05; **p < 0.005; NS, not significant.

To evaluate the significance of ADAM17 ECD positioning on the iRhom2 IHD, we introduced point mutations at the contact points (Figure 4A Site 1 and Site 2) in the ADAM17 ECD and assessed their effects on ADAM17 maturation and the stimulated shedding of AP-TGFα using the ADAM17 activator, phorbol 12-myristate 13-acetate (PMA) ^20, 62^. Among the tested mutations, R625E at Site 2 caused a significant decrease in AP-TGFα processing upon stimulation (Figure 4C). In contrast, the R610E mutation that is outside the iRhom2 IHD binding interface, did not affect AP-TGFα processing and functioned similarly to wild-type ADAM17. Despite the D and C domains of ADAM17 having few secondary structural features, they are stabilized and rigidified by a network of 13 disulfide bonds. Notably, the N480E mutation in site 1, located approximately 40 Å from R625 on the opposite side of the iRhom2 IHD interface, completely abolished AP-TGFα-stimulated shedding (Figure 4C). All mutant ADAM17 proteins still underwent proper maturation, a process dependent on iRhom binding (Supplementary Figure 4D). These findings support a model where the ancillary domains of ADAM17 serve as a rigid scaffold, stabilized by the iRhom IHD, that controls the activity of ADAM17, perhaps by allosteric control of the activity of the M domain or alignment with the cleavage site of its substrate.

Conservation analysis of ADAM17 revealed strong preservation in the catalytic cleft and a continuous interface extending down its transmembrane α-helix, suggesting an evolutionarily conserved functional significance for these residues (Figure 4 Site 3 and Supplementary Figure 4A-B). While iRhom2 exhibits low overall surface conservation, strong conservation is observed in TM1 and portions of the IHD hinge that interact with the ADAM17 TM α-helix (Supplementary Figure 4B). To explore the role of this conserved TM interface, point mutations were introduced into the ADAM17 transmembrane domain, and the stimulated shedding of AP-TGFα was evaluated (Figure 4C). Individual mutations of the ADAM17 TM attenuated AP-TGFα shedding from the cell surface with V673F, S680F and S688 having the most significant effect (Figure 4C). In contrast, a mutation (V678F) located on the opposite face of the ADAM17 TM α-helix relative to iRhom2 interaction, showed no defects in stimulated shedding of AP-TGFα. These findings emphasize the critical role of the transmembrane interface in binding ADAM17 and facilitating its functional regulation as part of the complex.

### Prodomain Removal is a Prerequisite for ADAM17 Sheddase Activity

The conversion of zymogen ADAM17 to its functionally mature state proceeds through the proteolytic processing by pro-protein convertases at two locations within the prodomain: the first at residue R58, designated the “upstream” processing site (US), and the second at the boundary site (BS) between the pro- and M domains ^24, 25^ (Figure 1A and 2A). In our cryo-EM structures, the BS and preceding region leading to the inhibitory cysteine-switch are missing in the density map, presumably due to high flexibility imposed by the binding of the MEDI3622. However, the US is positioned proximal to and is stabilized by the iRhom2 HD (Figure 1A). The stabilization of the US loop is underscored by its absence in the isolated cryo-EM structure of the MEDI3622 F_ab_-ProM domain (Supplementary Figure 2B).

To investigate the role of prodomain removal in ADAM17 activation, mutations were introduced at the US site (R58A) and the BS (R211A/R214G), in order to ablate furin processing and prevent ADAM17 maturation (Figure 5A). The impact of these mutations on ADAM17 proteolysis and maturation were assessed using the AP-TGFα shedding assay (Figure 5B-C). We found that mutation of the US site, either alone or combined with the boundary site, had the same effect on as the inactivating catalytic site mutant (E406A) on the basal and stimulated shedding of TGFα from the cell surface (Figure 5C). Mutating the BS site only reduced, but did not abolish TGFα shedding. Finally, expression of a mature form of ADAM17 lacking the entire prodomain (ΔPro) failed to elicit TGFα shedding (Figure 5C). These findings indicate that the presence of the prodomain during biosynthesis and cleavage at the US site is required for the stimulated shedding by ADAM17 in cells

**Figure 5:**
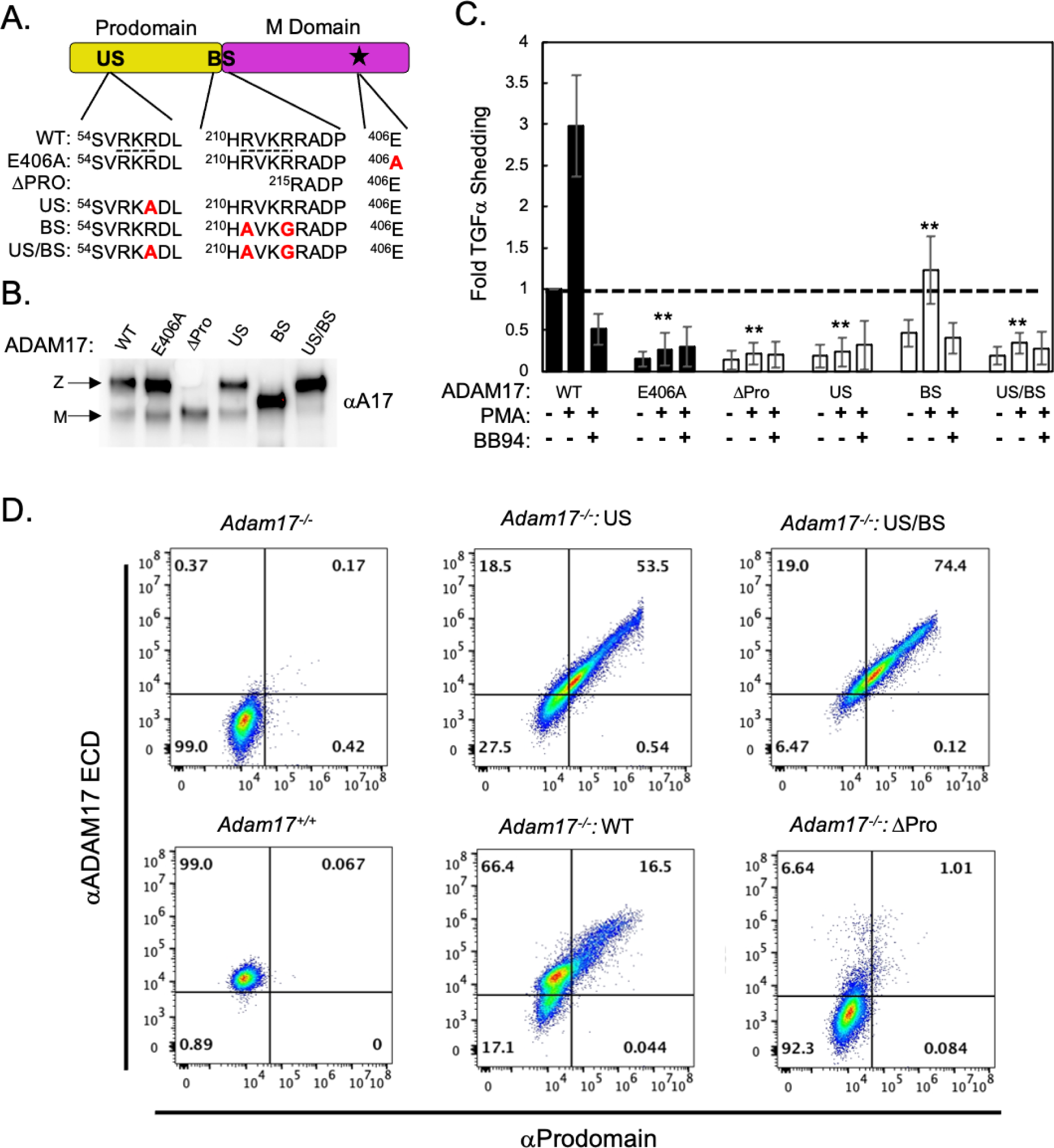
Analysis of Prodomain Processing in ADAM17 Activity. **(A)** Domain organization of the ADAM17 Prodomain and M Domain, highlighting the positions of the US, BS, and active site (depicted as a star), with the sequence alignments for the expression constructs. The consensus proprotein convertase processing sites are underlined with the point mutations to block maturation and enzyme activity indicated in red. **(B)** Western blot analysis of ConA-enriched lysates from cells transfected with ADAM17 variants: catalytically inactive E406A, ΔProdomain, US, BS, and US/BS mutations. To further distinguish between the zymogen (Z) and mature (M) forms of ADAM17, indicated with arrows, samples were treated with PNGasF. **(C)** Stimulated shedding assay of TGFα mediated by ADAM17. *adam17^−/−^* U2OS cells were co-transfected with AP-TGFα and either WT ADAM17 or ADAM17 prodomain mutants, as indicated. Shedding assays were conducted in the presence and absence of the pan-metalloprotease inhibitor BB94 to measure stimulated shedding activity. Data were normalized to the unstimulated WT ADAM17 control, in the absence of BB94. Error bars represent the mean ± SD from *N = 4* independent experiments. Statistical significance of PMA-stimulated mutant samples compared to WT was assessed using an unpaired, two-tailed t-test: *p < 0.05; **p < 0.005; NS, not significant. **(D)** Flow cytometry analysis of cell surface ADAM17 using selective antibodies targeting the prodomain and ectodomain. WT U2OS cells were compared to *adam17^−/−^* U2OS cells transfected with either WT ADAM17 or maturation-deficient ADAM17 mutants, including US/BS, US, and ΔProdomain. Quadrant gates were established based on the fluorescence signals from *adam17^−/−^* U2OS cells, which served as the negative control.

To further investigate the mechanistic defects caused by the US mutation, we used flow cytometry to detect the levels of the ECD and pro-domains of WT endogenous ADAM17 and overexpressed WT and US mutant forms at the cell surface. Surface staining of the US and US/BS ADAM17 mutants revealed strong correlation between cells stained with both an ADAM17 ECD antibody and an ADAM17 prodomain antibody (Figure 5D). Notably, expressing wild-type ADAM17 in ADAM17-null cells revealed that only cells with the highest ADAM17 ECD staining were double-stained with the prodomain antibody. Furthermore, cells expressing endogenous ADAM17 exhibited lower surface levels of ADAM17 compared to WT-transfected cells, with no prodomain detected on the cell surface. Moreover, the ΔPro ADAM17 mutant showed little surface staining of ADAM17 ECD, suggesting that it suffered from significant trafficking defects to the cell surface (Figure 5D). Taken together, this analysis supports a molecular model in which the prodomain is required for proper folding and exit of the iRhom2-ADAM17 complex from the ER. Pro-ADAM17 undergoes processing by pro-protein convertases, such as furin, in the trans-Golgi network, resulting in dissociation of the pro-domain from mature ADAM17 prior to reaching the cell surface.

## DISCUSSION

Here, we present the cryo-EM structure of the zymogen ADAM17 bound to iRhom2. The structure reveals critical interactions between the transmembrane α-helices and extracellular domains of both proteins. Moreover, it highlights a central role for the cytoplasmic “re-entry” loop in iRhom2 as a regulator of ADAM17 function, which was further corroborated by cell-based studies. Overall, our findings offer detailed new insights into the regulatory mechanisms controlling ADAM17 maturation and activity.

In our investigation, we found that the iRhom2 cytoplasmic region was a key feature in regulating enzyme function and in generating the zymogen ADAM17/iRhom2 complex that were amenable for structure determination. Interestingly, patients with gain-of-function mutations in a conserved cytoplasmic sequence in iRhom2 suffer from Tylosis with Oesophageal cancer (TOC) ^63^. Moreover, mice carrying deletions in this conserved region of the cytoplasmic domain exhibit dysfunctional ADAM17 activity and curly hair and bare skin ^45, 64, 65, 66^. The iRhom cytoplasmic domain undergoes post-translational phosphorylation, which modifies its intracellular interactions with regulatory binding proteins. Notably, the FERM domain-containing protein 8 (FRMD8, also known as iTAP) and 14-3-3 adaptor proteins have emerged as key regulators of ADAM17 activity, influencing the half-life and activity of mature ADAM17 ^40, 41, 43, 44^. In macrophages lacking iRhom2, or in other tissues devoid of both iRhom1 and -2, there is no detectable mature ADAM17, although a stable pro-form of ADAM17 is present, likely residing in the ER ^32, 33, 34, 35^. Conversely, in the absence of ADAM17, iRhom2 stability is significantly compromised, suggesting that iRhom2’s stability hinges on proper assembly with ADAM17 ^48^. Supporting this, FRMD8 knockout mice exhibit minimal mature ADAM17, aligning with the notion that FRMD8 is crucial for the proper assembly of the iRhom2/ADAM17 complex, thereby enabling ER exit ^67^. Additionally, Lu *et al.* have reported that co-expression of FRMD8 was vital for generating recombinant protein suitable for their structural studies of iRhom2/ADAM17 ^49^. In our investigation, we found that removing part of the iRhom2 cytoplasmic domain, including the FRMD8-binding site ^41, 44, 67^ was a crucial step in producing a stable ADAM17-iRhom2 complex that was amenable to structural characterization. FRMD8 binding to the full length iRhom2 cytoplasmic domain may mirror the effects observed upon deletion of portions of the iRhom2 cytoplasmic domain, including the FRMD8 binding site. Binding of FRMD8 or deletion of its binding site potentially facilitates the proper assembly and folding of the iRhom2/ADAM17 complex in the endoplasmic reticulum (ER), which appears to be a prerequisite for ADAM17’s exit from the ER.

We and others have shown that proper ADAM17 trafficking and function require initial translation as a zymogen with an intact prodomain, followed by maturation involving the cleavage and removal of the prodomain in the trans-Golgi network ^24, 54, 55^. The importance of the prodomain is underpinned by its central position within the zymogen ADAM17-iRhom2 cryo-EM structure, which shows that the prodomain bridges the ADAM17 M domain to the iRhom2 IHD. Indeed, interactions between zymogen ADAM17 and iRhom2 have been observed to be more stable than those of the mature ADAM17-iRhom2 complex ^33^. As this manuscript was in preparation, a cryo-EM structure of the zymogen ADAM17-iRhom2 complex was published that readily superimposes (RMSD = 0.813 Å) onto the structure presented here ^49^. Lu et al. proposed that the release of the processed prodomain represents the mechanism underlying ADAM17 activation by stimuli, such as the phorbol ester PMA. However, this interpretation is based on the properties of a mutant ADAM17 in which the US site cannot be processed, similar to the construct described above and to a previously reported construct that does not exhibit sheddase activity towards TGFα ^25^. Here, we confirm that mutating the US site severely attenuates ADAM17 sheddase activity. Furthermore, we show that the fully processed prodomain does not remain associated with mature ADAM17, suggesting that it is rapidly removed following processing by a pro-protein convertase in the trans-Golgi network. Thus, the prodomain cannot be detected on the surface of unstimulated wild-type cells and is unlikely to regulate ADAM17’s rapid activation in response to stimuli. Instead, we propose that ADAM17 activation depends on allosteric conformational changes in the iRhom2-ADAM17 complex that are likely controlled by the iRhom2 re-entry loop.

Comparative analysis of the ADAM17-iRhom2 and ADAM10-tspan15 complexes suggests that these multi-transmembrane accessory proteins serve to tether the ancillary ADAM domains to the cell surface, potentially restricting the conformational states of the ADAM extracellular region. Our structure and mutational analysis support a model in which the M domain is positioned by the IHD to promote enzyme function. In agreement, the Dusterhoft group showed that a W567S mutation, buried in the iRhom2 IHD, likely compromises the IHD’s structural integrity and its binding to the ADAM17 ECD, thereby attenuating ADAM17 function in cells without disrupting complex formation with iRhom2 ^68^. Moreover, we show that key residues (R625, K626, and K628) interact with an oppositely charged surface on the iRhom2 HD, positioning the ADAM17 ECD relative to the cell surface. It is tempting to speculate that binding proteins for both ADAM10 and ADAM17 serve to position the rigid C+D domains, thus aiding in the precise positioning of the M domain. This restriction could function akin to a molecular ruler, as described for ADAM10 ^52^, or as a conformational scaffold for the catalytic domain, as suggested for the iRhom2-ADAM17 complex ^69^, contributing to their substrate selectivity. This suggests that therapeutic targeting of the ADAM17-iRhom2 ECD interface, particularly the iRhom2 HD, could be a viable strategy for inhibiting ADAM17 activity. This approach may create more selective inhibitors than antibodies targeting the ADAM17 protease directly, as anti-iRhom2 antibodies would not interfere with iRhom1-ADAM17 complexes. This selectivity could be especially relevant in chronic inflammatory diseases such as rheumatoid arthritis and systemic lupus erythematosus glomerulonephritis, where dysregulated iRhom2-ADAM17 activity plays a critical role ^5, 6^.

Stimulated ADAM17 activity is dependent on complex formation with iRhom2, which requires the TM helices and specific regions of the iRhom2 cytoplasmic domain ^28, 40, 41, 43, 44, 46^. The TM regions of ADAM17 and iRhom2 form a conserved interaction site that regulates ADAM17’s initial assembly and activity. Swapping ADAM17’s TM with that of an unrelated type-1 TM α-helix from BTC or CD62L abolishes stimulated activity, whereas restoring ADAM17-specific TM residues reinstates activity ^47^. Prior studies have shown that the cytoplasmic domain of ADAM17 is dispensable for rapid activation by various stimuli, positioning iRhom’s as key regulators of ADAM17 activity ^26, 28^. Our finding that a conserved cytoplasmic re-entry loop near TM1 affects ADAM17 activity provides a possible mechanistic explanation. Constructs of iRhom2 with nearly complete cytoplasmic deletion but retention of the re-entry loop (Δ363) activate ADAM17-dependent shedding without stimulation. This activation is lost with mutations in the re-entry loop near TM2 (R367E) or deletion of the re-entry loop (Δ381). Intriguingly, the re-entry loop within the core rhomboid fold of iRhom2 is positioned between TM2 and TM5, which in rhomboid protease proteins typically create a substrate-binding pocket guiding substrates to the active site. In contrast, iRhom2 binds ADAM17 primarily along the outward-facing side of TM1, leaving the TM2-TM5 interface unobstructed. TM2’s functional relevance is further underscored by missense mutations in mouse iRhom2 TM2 (V645E and L655Q) that impair the proteolytic processing of TNFR2 and TNFα ^70, 71^. Furthermore, deleting nine residues in the iRhom2 IHD near TM2 abolishes iRhom2/ADAM17 activity in iRhom1/2^−/−^ knockout cells ^47^. Notably, a mutation in the TM2 of iRhom1 (G665W, aligning with iRhom2 G666) has been linked to cardiomyopathy in a patient ^72^. Collectively, these results indicate that the re-entry loop is crucial for regulating ADAM17 activity by iRhom2, likely by affecting the relative positions of TM2 and TM5. We speculate that the TM2 may act as a fulcrum, transmitting positional changes across the membrane to affect the orientation of the extracellular iRhom2 HD, which resides between TM1 and TM2. Furthermore, changes in the iRhom2 HD could, in turn, modulate ADAM17 function, explaining why ADAM17’s cytoplasmic domain is dispensable for activation while its TM domain is essential.

### Limitations of the Study

Despite these advances, certain limitations remain. Our structure lacks the cytoplasmic regulatory elements of iRhom2, including the FRMD8 binding sites, that are involved in controlling stimulated shedding. Future studies will be required to explore the possibility of distinct active and inactive conformations of the ADAM17-iRhom2 complex. Furthermore, the molecular and atomic details underlying ADAM17 substrate processing and protein substrate binding and selectivity are still not fully understood. Moreover, the details of how exactly stimulation from inside the cell results in changes to ADAM17 activity in the extracellular space remain to be established. Finally, the conformation of ADAM17 bound to iRhom1 remains to be established by Cryo-EM, which would address whether these proteins might direct ADAM17 into different conformational states. Future research should aim to resolve these gaps, focusing on the regulatory mechanisms of iRhoms and the active states of the ADAM17-iRhom2 complex. Understanding these aspects could significantly enhance our ability to develop targeted therapies for diseases associated with dysregulated ADAM17 activity.

## METHODS AND MATERIALS

### Cell Lines

All cell lines used in this study (U2OS, Expi293, and SF9) have been described previously and were maintained in their respective media ^52, 59^. The generation of the *Adam17^⁻/⁻^* U2OS cell line was performed by the Cincinnati Children’s Hospital Medical Center Transgenic Animal and Genome Editing Core Facility, which introduced a nonsense mutation into exon 2 of the *Adam17* gene locus. Successful genome editing by CRISPR-Cas9 repair was validated through sequencing, and western blot analysis confirmed a complete loss of ADAM17 protein expression.

### Expression Constructs

The ADAM17 expression vectors were subcloned from pRK5M-ADAM17myc (Addgene: Plasmid #31714) into the pEG BacMam expression vector (Addgene: Plasmid #160451) ^74, 75^. The extracellular ADAM17 pro- and metalloprotease domains (residues 1– 477) were cloned into the pLib vector (Addgene: Plasmid #80610), which includes a 6x HisTag at the carboxyl terminus ^76^. All iRhom2 expression constructs were PCR-amplified from a cDNA (Origene) and sequentially subcloned into the p-mVenusC1 vector (Addgene: Plasmid #27794), followed by cloning into pEG BacMam as mVenus-iRhom2 chimeric proteins ^77^. The MEDI3622 antibody F_ab_ fragment sequence was obtained from World Intellectual Property Organization Patent #WO 2016/089888 A1. Geneblock fragments encoding the light and heavy chains were cloned into the expression vectors pD2610-v5 (ATUM) and pFUSE-hIgG1-Fc1 (Invitrogen), respectively, with the latter containing a 3C protease site upstream of the Fc region. The AP-TGFα reporter construct was generated by cloning a Geneblock encoding TGFα (residues 40–161) into the pTAG5-AP vector (GenHunter: Cat#QV5). All point mutations in ADAM17 and iRhom2 were introduced using overlapping PCR primers. The ΔPro ADAM17 (amino acid 215-824) was inserted into the pRK5M-myc expression vector behind its native signal sequence using Infusion Cloning (Takara Bio Inc.). Each expression construct was validated by Sanger sequencing.

### Recombinant Protein Production and Isolation

The heavy and light chains of the MEDI3622 antibody were co-transfected (1 μg/mL, 1:1 Light to Heavy Chain ratio) into Expi293F cells at a density of 2 × 10⁶ cells/mL using a 1:1 DNA to 40 kDa polyethyleneimine HCl MAX ratio. After 24 hours, the cell culture was supplemented with 450 mg/mL of D-(+)-glucose and 5 mM valproic acid sodium salt. Transfected cultures were incubated at 37°C while shaking for a total of five days. Conditioned media was collected by centrifugation at 4000 rpm for 30 minutes and supplemented with 20 mM HEPES, pH 8.0, and 150 mM NaCl. The clarified media was applied to a Protein-A Sepharose column, extensively washed with 20 mM HEPES, pH 8.0, and 150 mM NaCl, and the bound protein was eluted in a 40 mM glycine, pH 3.0, and 150 mM NaCl buffer. The eluate was immediately neutralized with 1 M HEPES, pH 8.0. The MEDI3622 antibody was treated overnight at 4°C with 100 ng of 3C protease per 5 μg of antibody to release the F_ab_ fragment from the Fc region. The MEDI3622 F_ab_ fragment was purified using a Superdex 200 Increase (Cytiva) column equilibrated with 20 mM HEPES, pH 8.0, and 150 mM NaCl.

The A17 ProM Domain protein, containing the single point mutations N174Q, R58A, R211A and R214A, was expressed from baculovirus-infected *Spodoptera frugiperda*-derived ovarian cells (SF9) at a density of 3.0 × 10⁶ cells/mL. Cultures were incubated at 27°C with shaking at 120 RPM for an additional four days. Conditioned media was collected by centrifugation and supplemented with 20 mM Tris buffer, pH 7.5, containing 150 mM NaCl, 5 mM CaCl₂, 1 mM NiCl₂, and 0.01 mM ZnCl₂. The media was clarified by centrifugation to remove residual debris and applied to a Ni-NTA column. The column was extensively washed with buffer containing 20 mM HEPES, pH 8.0, 100 mM NaCl, and 20 mM Imidazole and bound protein was eluted in the same buffer with 250 mM imidazole. Fractions were concentrated with centrifugal filters and buffer exchanged to remove excess salt and applied to an anion exchange and eluted over a 1 – 50% gradient of 20 mM HEPES, pH 8.0, 1000 mM NaCl. Fractions containing the ADAM17 ProM domains were concentrated and mixed with excess MEDI3622 F_ab_ and isolated to homogeneity with a Superdex 200 10/300 Increase (Cytiva) size exclusion chromatography column in 20 mM HEPES, pH 8.0, and 150 mM NaCl for use in cryo-EM sample preparation.

Full-length ADAM17 (E306A) and mVenus-iRhom2 (residues 364–824) proteins were co-expressed in baculovirus-infected Expi293F cells at a volumetric ratio of 1:3, corresponding to 6% v/v of the total cell volume. After 24 hours, the infected Expi293F cultures were supplemented with 450 mg/mL D-(+)-glucose and 5 mM valproic acid sodium salt. The cells were harvested after 72 hours by centrifugation at 4,000 rpm for 30 minutes and resuspended in a homogenization buffer containing 20 mM HEPES at pH 8.0, 300 mM NaCl, 1% lauryl maltose neopentyl glycol (LMNG) (Anatrace), 0.1% cholesteryl hemisuccinate (CHS) (Anatrace), 1 mM CaCl₂, and 30% glycerol. The suspension was homogenized by stirring at 4°C for one hour. Cellular debris was removed by centrifugation at 14,000 rpm for 45 minutes, and the resulting supernatant was filtered through a 0.7 μm glass microfiber filter (Watman). The clarified supernatant was applied to a CNBr-GFP Nanobody affinity column and extensively washed with two column volumes of a buffer containing 20 mM HEPES at pH 8.0, 300 mM NaCl, 0.1% LMNG, 0.01% CHS, 0.1 mM CaCl₂, and 3% glycerol. To exchange detergents, the bound protein was sequentially washed with an exchange buffer containing 20 mM HEPES at pH 8.0, 150 mM NaCl, and 0.1% glycol-diosgenin (GDN) (Anatrace), followed by a second buffer containing 20 mM HEPES at pH 8.0, 150 mM NaCl, and 0.05% GDN. The protein was eluted using a buffer composed of 100 mM glycine at pH 3.0, 150 mM NaCl, and 0.05% GDN and was immediately neutralized with 10 mL of a neutralizing buffer containing 450 mM HEPES at pH 8.0, 150 mM NaCl, and 0.05% GDN. The eluted protein was concentrated using a 100 kDa centrifugal filter at 4,000 rpm until the total volume was reduced to 1 mL. The sample zymogen ADAM17-iRhom2 complex was mixed with MEDI3622 F_ab_ then applied to a Superose 6 Increase 10/300 GL (Cytiva) size-exclusion chromatography column, and fractions were analyzed by SDS-PAGE analysis and pooled based on sample homogeneity. The pooled fractions were further concentrated and immediately used for cryo-EM sample preparation.

### Cryo-EM Sample Preparation and Data Acquisition

The ADAM17 – iRhom2 and ProM domain in complex with the MEDI3622 F_ab_ were concentrated to 5.3 mg/mL and 0.25 mg/mL, respectably. These samples were individually applied to AltrAuFoil® R 1.2/1.3 grids that were glow discharged using a PELCO easiGlow™ Glow Discharge Cleaning System; 0.39 mBar, 20 mA, glow time/hold 30/10s. Grids were blotted with a force between 1-10, for times ranging 1-11s and plunge frozen in liquid ethane using a Vitrobot Mark IV System.

ADAM17 – iRhom2 grids were imaged at the Vanderbilt School of Medicine Center for Structural Biology on a FEI Titan Krios operated at 300 kV with a K3 BioQuantum direct electron detector camera in counting mode. Data sets were collected at a nominal magnification of 105,000x with a pixel size of 0.822 Å and defocus range between −0.4 to −2.2 µm. Two data sets were collected using a total dose of 62.2 e^−^/Å^2^ and 56.3 e^−^/Å^2^ were merged, totaling 24,828 movies, that were used for single particle analysis structure determination. The ADAM17 Pro-M Domain grids were imaged at the University of Cincinnati School of Medicine Center for Advanced Structural Biology on a Glacios operated at 200 kV with a Falcon 4D direct electron detector. A single data set, totaling 8,704 movies, was collected at a nominal magnification 165,000x with a pixel size of 0.69 Å, defocus range −0.4 to −2.0 µm and total dose of 40.0 e^−^/Å^2^ was used for structure determination.

### Cryo-EM Single Particle Analysis and Model Building

All cryo-EM data were processed using CryoSPARC on the GPU cluster maintained by the University of Cincinnati Advanced Research Computing Center and all cryo-EM software, excluding CryoSparc, was maintained in collaboration with SBGrid ^78^. Movies were motion-corrected using patch motion correction, and contrast transfer function (CTF) parameters were estimated with patch CTF estimation. Movies were curated based on total full-frame motion distance (below 40 pixels), relative ice thickness (below 1.1), and CTF fit resolution (below 4.5 Å). For the ProM domain and ADAM17–iRhom2 structures, this process resulted in 5,673 and 18,680 movies, respectively, being selected for structure determination.

Initial particles were picked using a Laplacian of Gaussian autopicking approach on a subset of 300 curated movies to generate template 2D classes for particle picking across a larger dataset. For the ADAM17–iRhom2 complex, these 2D templates were used to pick an additional 2,084,443 particles from 5,652 curated movies. Iterative 2D classification reduced this set to 440,135 particles, which were used for iterative *ab initio* reconstruction, yielding an initial 6 Å density map for the ADAM17–iRhom2 complex. This particle stack served as a reference for training the Topaz particle-picking model, which subsequently identified 2,965,292 particles from the total curated movie set ^79^. After further curation through 2D classification, 2,421,726 particles remained. These particles underwent heterogeneous refinement using three input models, resulting in a 6.25 Å density map containing 250,122 particles. Iterative 3D classifications and local refinements improved the resolution to 3.53 Å. The final map was post-processed using DeepEMhancer with the high-resolution model ^80^. The overall workflow is depicted in Supplemental Figure 1B. AlphaFold predictive models for the iRhom2 – ADAM17 zymogen complex and MEDI3622 F_ab_ were fit into the density using ChimeraX ^81, 82^. Due to the slight structural reorganization of ADAM17 domains from the predictive AlphaFold model, the ADAM17 model was divided into three parts for fitting: the ProD domain, C + D domains and TM region. The model was built in Coot and refined in Phenix Real-Space Refine ^83, 84^. Model to map FSC was produced using Phenix Comprehensive Validation.

For the ProM domain, initial 2D classes were curated for use within the Topaz CryoSPARC workflow, resulting in 2,357,801 particles, which were reduced to 334,801 particles after 2D classification. This curated particle stack was used to generate three *ab initio* models, one of which represented the complete ProM-F_ab_ complex. These initial models underwent heterogeneous refinement against the 2,357,706-particle stack, followed by additional heterogeneous refinements to sort particles into the full complex map. The final density map was generated using a 3D-sorted particle set of 346,939 particles. Iterative *ab initio* modeling of two low-similarity classes (similarity score = 0.01), followed by NU-refinement and local refinement, resulted in a final density map with an overall resolution of 3.5 Å using 98,475 particles. The map was also post-processed using DeepEMhancer with the high-resolution model and the overall workflow is depicted in Supplemental Figure 2. The coordinates for the ProM domain and MEDI3622 F_ab_ from the complex described above were used to fit into the final density map using ChimeraX. This model was also built in Coot and refined in Phenix Real-Space Refine. Model to map FSC was produced using Phenix Comprehensive Validation.

### ADAM17 Ectodomain shedding Assay

The methods for ADAM17 basal and stimulated ectodomain shedding assays were adapted from previously established protocols with slight modifications ^39, 60^. In brief, wild-type (WT) or *Adam17^−/−^* U2OS cells were seeded into 6-well tissue culture plates and co-transfected with either mVenus, ADAM17, or iRhom2 expression constructs along with the AP-TGFα reporter at a 1:2 ratio using PEI MAX. For stimulated shedding assays, 24 hours post-transfection, cells were washed with DMEM and serum-starved for 1 hr. Following this, the culture media was replaced with media supplemented with either 0.6% DMSO, 25 ng/μL PMA, or 5 μM BB94 + 25 ng/μL PMA. After 1 hr, conditioned media and cell lysates were collected, and the amount of AP activity in the supernatant and lysates was measured at an absorbance of 405 nm over a 90-minute time course following the addition of 1 mg/mL 4-nitrophenyl phosphate (New England Biolabs), using a BioTek SynergyH1 microplate reader. Ratios of AP activity in the supernatant to the combined cell lysate and supernatant were calculated. Each sample was normalized to its DMSO control to assess stimulation changes in response to the different pharmacological treatments. For basal shedding assays, 24 hr post-transfection, cells were washed, and 1 mL of DMEM supplemented with either 0.6% DMSO or 5 μM BB94 was added. After 24 hr, AP activity and ratios were calculated as described for the stimulated shedding assay. All sample ratios were normalized to the mVenus basal shedding.

### Immunoprecipitation and Western Blotting

Expi293F cells were grown to a density of 1×10⁶ cells/mL and co-transfected with 1.5 μg of wild-type or mutant ADAM17 and 0.5 μg of mVenus-iRhom2 cDNA using Lipofectamine™ 2000 (ThermoFisher Scientific). After 24 hrs, cells were harvested by centrifugation at 14,000 RPM for 5 minutes and suspended in homogenization buffer containing 20 mM HEPES pH 8.0, 300 mM NaCl, 1% LMNG, 0.1% CHS, 1 mM CaCl₂, and 30% glycerol. The suspension was incubated for 1 hour. Lysates were clarified by centrifugation at 14,000 RPM for 5 min, and proteins were immunoprecipitated using CNBr-GFP Nanobody resin for 1 hour. The immunoprecipitation resin was extensively washed with 10x bead volume of homogenization buffer nine times to remove non-specifically captured proteins. Bound proteins were eluted by boiling the resin in 2x Laemmli Sample Buffer (BioRad #1610737), supplemented with 50 mM β-mercaptoethanol, for 5 minutes. The immunoprecipitated samples were analyzed by western blot for the presence of ADAM17 and iRhom2-GFP using αTACE (Cell Signaling, cat #6978) and αGFP (BioRad, cat #AHP975) antibodies, respectively.

### Flow Cytometry

WT and *Adam17^−/−^* U2OS cells were seeded in 6-well tissue culture dishes and transfected with 0.5 ug of the ADAM17 expression constructs, using PEI. After 48 hrs, cells were dislodged from the plate by trypsinization and isolated by centrifugation at 500 g for 5 minutes. Cells were stained for ADAM17 ECD and ADAM17 Prodomain with primary antibodies, αADAM17-alexafluor647 (R&D Systems) and αprodomain ADAM17 ^24^, respectively, at a 1:100 dilution for 90 minutes at room temperature in PBS supplemented with 0.5% m:v Bovine Serum Albumin. Cells were then washed with 10x volume of PBS and incubated for 60 minutes at room temperature with αRabbit-AlexaFluor488 (ThermoFisher) for detection of the prodomain antibody. Cells were subsequently washed with and resuspended in PBS for flow cytometry analysis. Single stain controls used for the ECD-ADAM17-alexafluor 647 and prodomain-αRabbit-AlexaFluor488 were WT U20S cells and adam17^−/−^ U20S transfected with ADAM17US/BS mutations, respectively. Cell surface staining was assessed on a Cytek Aurora and analyzed by FlowJo software (v 10.10), measuring co-emission of 647 nm and 488 nm. For flow cytometry analysis, events were gated by FSC-SSC to remove debris and then restricted to single cells. Cells Quadrant gates for the purpose of quantification and depiction were generated with use of *Adam17^−/−^* null cells.

### Quantification and Statistical Analysis

Results of statistical analyses are found in the figure legends for Figures 3, 4, and 5. All calculations of significance were determined in Microsoft Excel software package using an unpaired t test (two-tailed). Data are reported as mean ± standard deviation. Significance was determined by a p value of < 0.05, and annotated as *p < 0.05 and **p < 0.005. Not significant is annotated with NS.

## DATA AVALABILITY

Structural data for the MEDI3622 F_ab_-ADAM17-iRhom2 and MEDI3622 F_ab_-ADAM17 Pro-M domain complexes we be made available on the RCSB. Additional materials can be provided upon request to T.C.M.S. For further details or to revisit the analyses presented, inquiries may be directed to the lead contact.

## ACKNOWLEDGEMENTS

The authors thank the staff at the Vanderbilt Structural Biology Center, University of Cincinnati Center for Advanced Structural Biology, and University of Cincinnati Advanced Research Computing Center for their assistance with data collection and maintaining the computational cluster for data processing. In addition, we would like to thank Dr. Igal Ifergan for assistance with the flow cytometry analysis. Special thanks to Drs. Stephen Blacklow, Rhett Kovall, and Xiaowei Hou for critical reading and feedback on this manuscript. Financial support was provided by the University of Cincinnati Innovative Pilot Grant (to T.C.M.S) and by National Institute of General Medical Sciences R35GM146830 (to T.C.M.S) and R35GM134907 (to C.P.B).

## CONTRIBUTIONS

T.C.M.S. conceived the project. J.J.M., C.S., C.P.B., and T.C.M.S. designed the experiments. Cell-based assays were performed and analyzed by J.J.M., C.S., H.F.A., and T.C.M.S. Recombinant protein production was performed by J.J.M., C.S., M.R., and B.G. Cryo-EM data acquisition was carried out by J.J.M., with data processing completed by J.J.M., C.S., and T.C.M.S. The manuscript was written by T.C.M.S., C.P.B., J.J.M., C.S. and input from all other authors.

## COMPETING INTERST

C.P.B. holds a patent for a methodology to identify agents that synergize with iRhom inhibitors. In partnership with the Hospital for Special Surgery, C.P.B. has developed iRhom2 inhibitors and co-founded SciRhom, a Munich-based start-up, to bring these inhibitors to market.

