## Supplemental Figures for "Structural Insights into the Activation and Inhibition of the ADAM17-iRhom2 Complex"

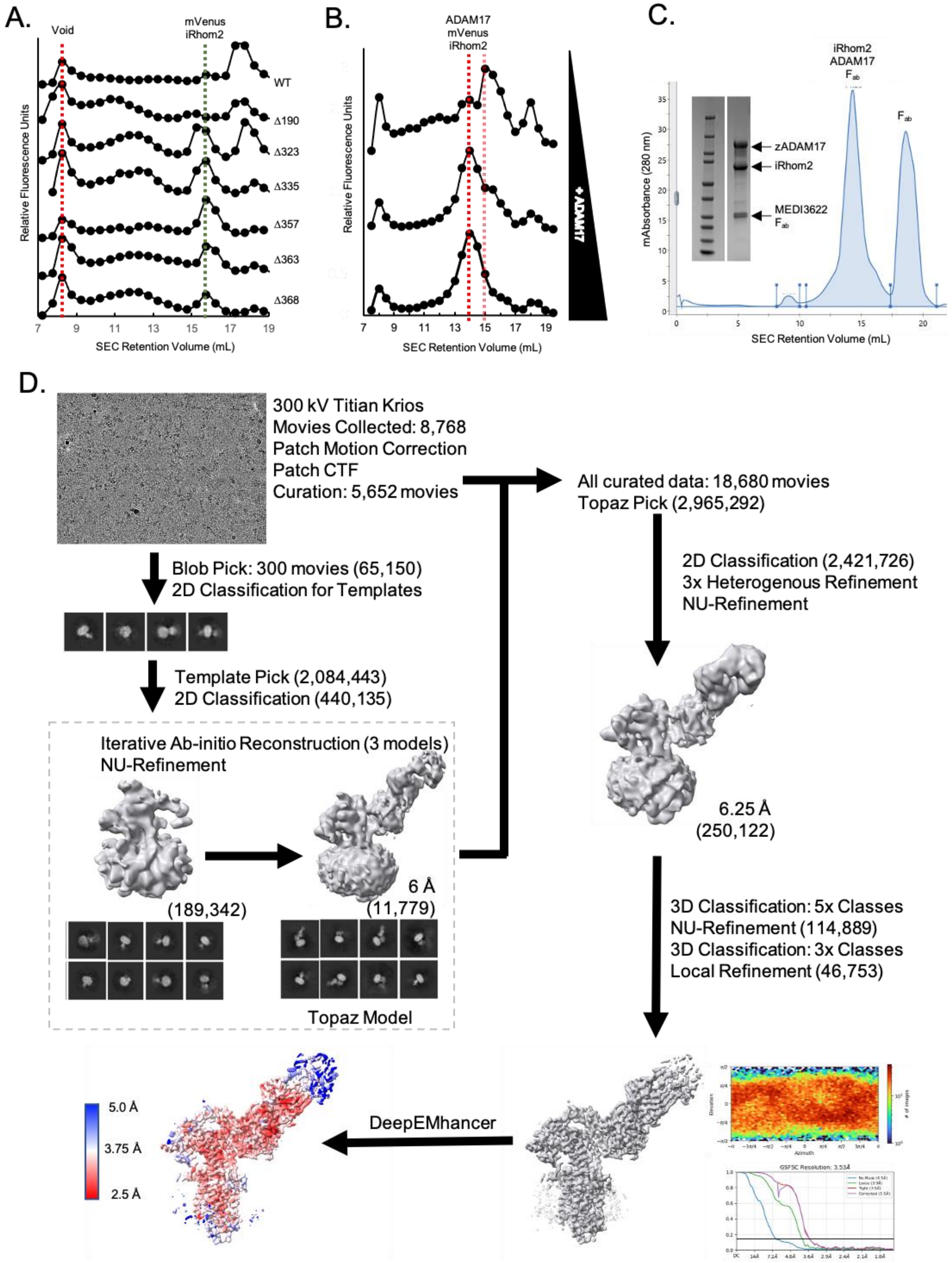

A.
Void
mVenus
iRhom2
WT
Δ190
Δ323
Δ335
Δ357
Δ363
Δ368
Relative Fluorescence Units
SEC Retention Volume (mL)
B.
ADAM17
mVenus
iRhom2
+ ADAM17
Relative Fluorescence Units
SEC Retention Volume (mL)
C.
iRhom2
ADAM17
Fab
Fab
zADAM17
iRhom2
MEDI3622
Fab
mAbsorbance (280 nm)
SEC Retention Volume (mL)
D.
300 kV Titian Krios
Movies Collected: 8,768
Patch Motion Correction
Patch CTF
Curation: 5,652 movies
Blob Pick: 300 movies (65,150)
2D Classification for Templates
Template Pick (2,084,443)
2D Classification (440,135)
Iterative Ab-initio Reconstruction (3 models)
NU-Refinement
(189,342)
6 Å
(11,779)
Topaz Model
All curated data: 18,680 movies
Topaz Pick (2,965,292)
2D Classification (2,421,726)
3x Heterogenous Refinement
NU-Refinement
6.25 Å
(250,122)
3D Classification: 5x Classes
NU-Refinement (114,889)
3D Classification: 3x Classes
Local Refinement (46,753)
DeepEMhancer
5.0 Å
3.75 Å
2.5 Å
GSFSC Resolution: 3.53Å

**Supplementary Figure 1: Structure Determination of the zymogen ADAM17-iRhom2 Workflow.** **(A)** Fluorescence-detection size exclusion chromatography (FSEC) of cytoplasmic truncated iRhom2 variants. The column void is indicated by the red line, and mVenus-iRhom2 is represented by the black line. **(B)** FSEC analysis of Δ363-iRhom2 co-expressed with increasing amounts of ADAM17. Peaks are annotated to indicate the formation of the ADAM17-iRhom2 complex with a leftward shift in the dashed red line. **(C)** Size exclusion chromatography (SEC) of the purified zymogen ADAM17-Δ363-iRhom2-MEDI3622 $F_{ab}$ complex. Annotated peaks correspond to components of the complex. *(Inset)* Coomassie-stained SDS-PAGE shows the purity of the zymogen ADAM17, Δ363-iRhom2, and MEDI3622 $F_{ab}$ complex used for cryo-EM sample preparation. **(D)** Workflow diagram illustrating the process used to determine the structure of the zymogen ADAM17-Δ363-iRhom2-MEDI3622 $F_{ab}$ complex.

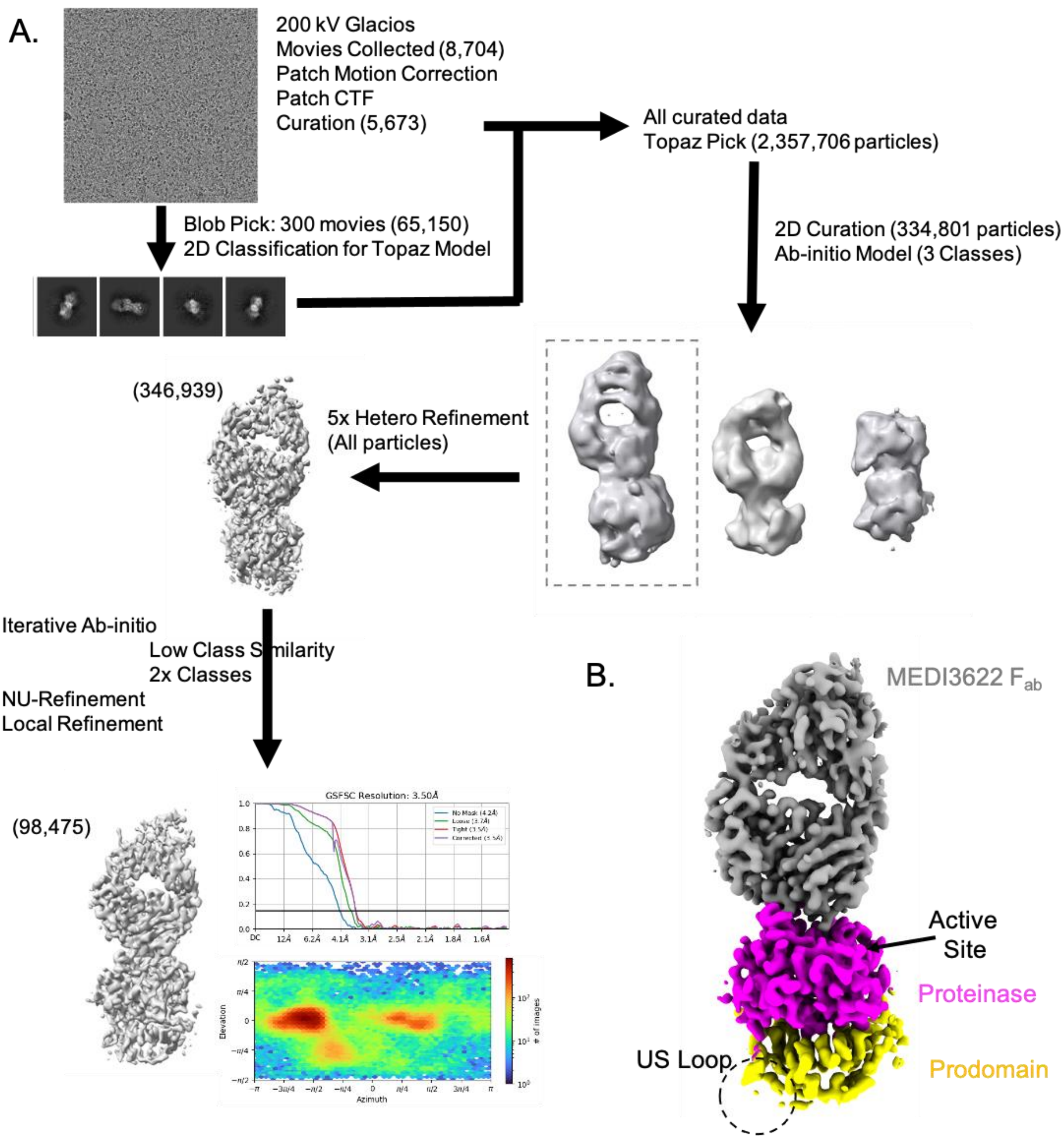


**Supplementary Figure 2: Structure Determination of the ADAM17 Pro-M Domain Workflow. (A)** Workflow diagram illustrating the process used to determine the structure of the Prodomain and Metalloproteinase domain of ADAM17 in complex with the MEDI3622 $F_{ab}$. (B) Final density map of the ADAM17 ProM-MEDI3622 $F_{ab}$ complex colored by domain as done in Figure 1.  The prodomain US proprotein convertase site location is designed with a dashed circle.

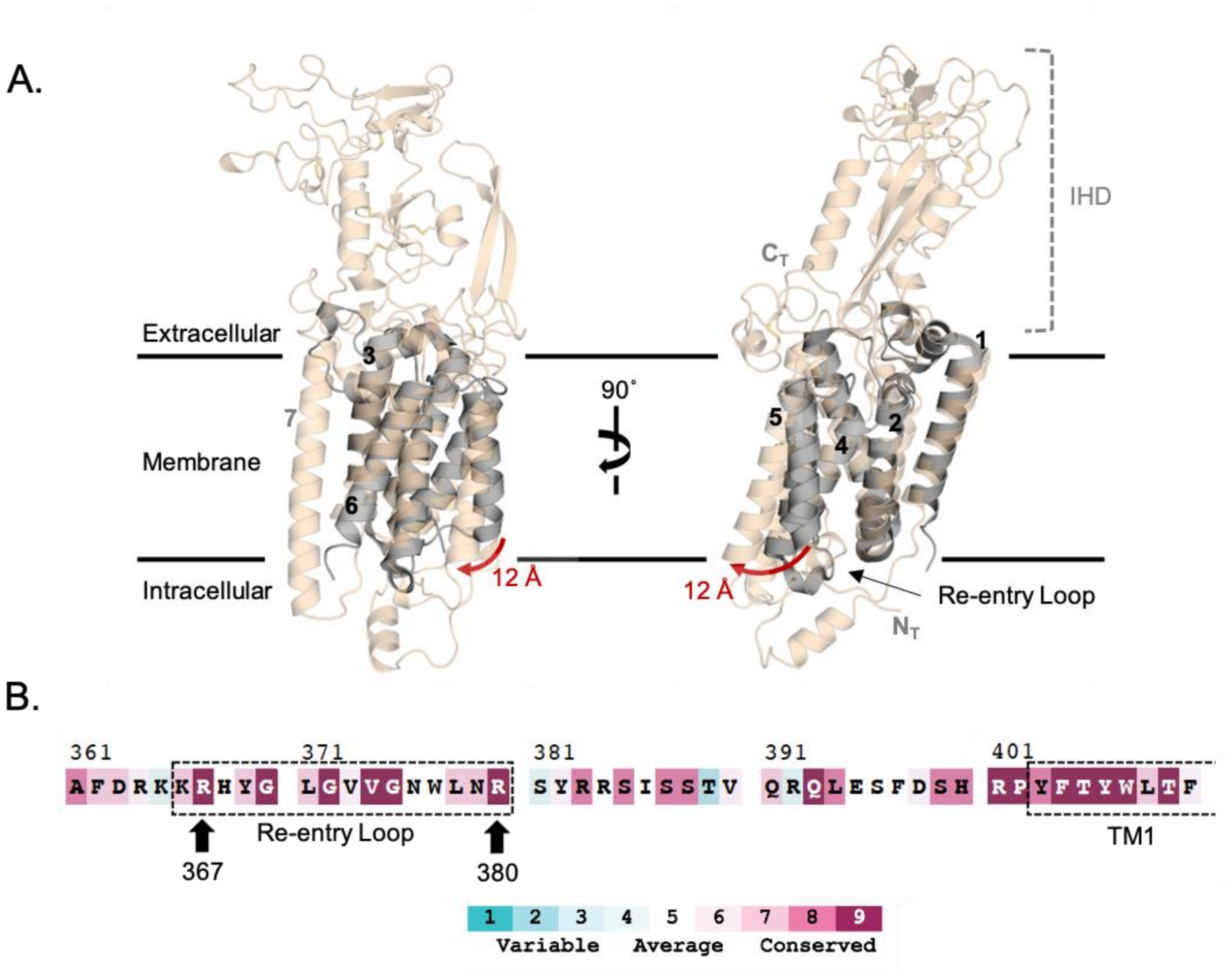


**Supplementary Figure 3: Structural Elements of iRhom2 in ADAM17 Substrate Recognition. (A)** Cartoon representation and structural superimposition of GlpG (dark gray; RCSB PDB: 2IC8) with iRhom2 (tan), shown embedded in the cell membrane. The re-entry loop of iRhom2 is annotated with an arrow, and the iRhom2 homology domain (IHD) is indicated with a dashed bracket. A red arrow highlights the displacement of iRhom2 TM5 relative to GlpG TM5 to accommodate the re-entry loop. **(B)** Conservation analysis of iRhom2 residues 361–410 using a color-coded scale from variable (teal) to conserved (maroon) [73]. The re-entry loop and TM1 regions are enclosed within a dashed box. Conserved Arg residues used in Figure 3D-E are annotated with arrows.

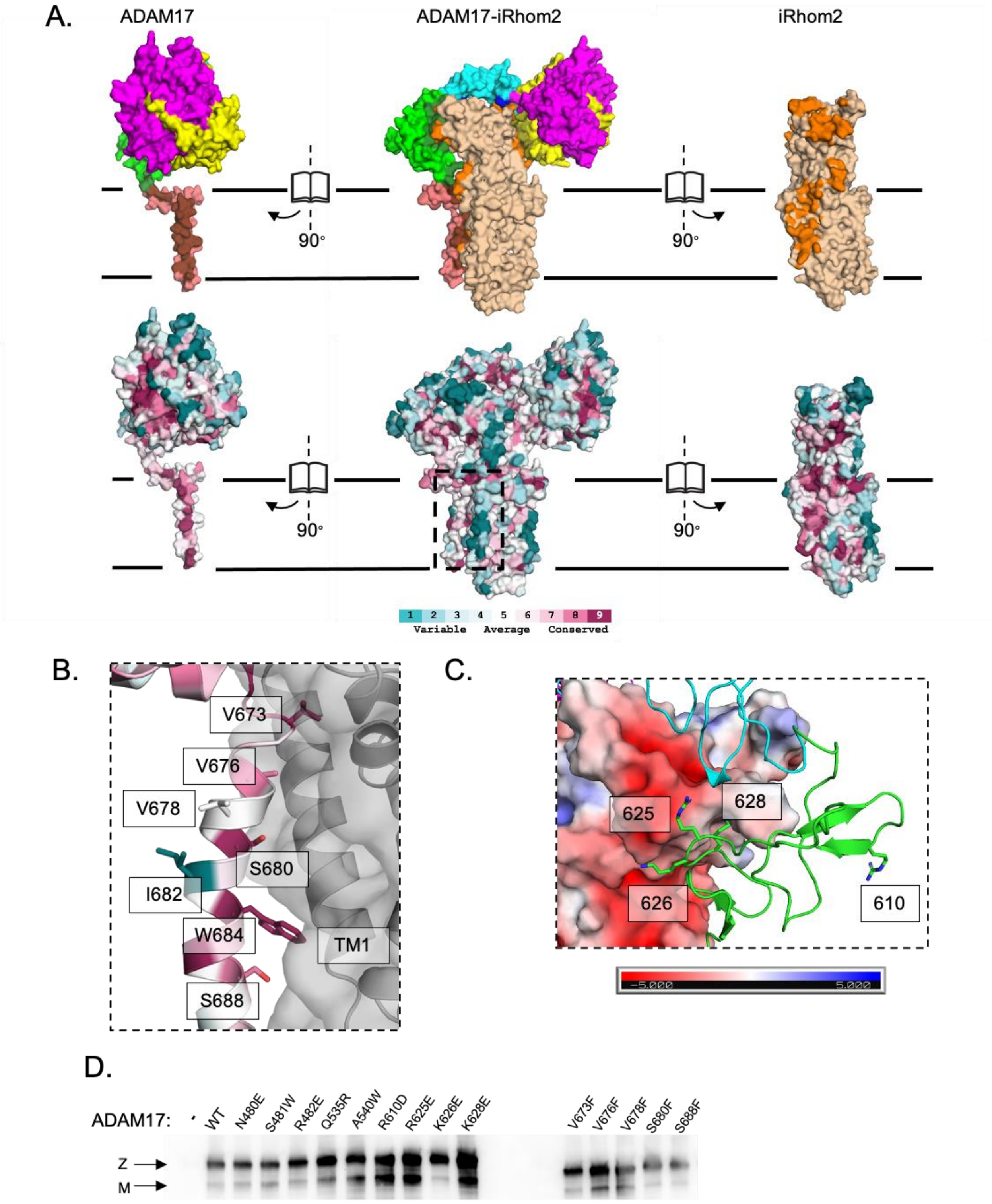


**Supplementary Figure 4: Conservation analysis of the zymogen ADAM17-iRhom2.**

**(A)** Surface representation of the zymogen ADAM17-iRhom2 complex, with domains colored as in Figure 1. Open-book views illustrate the contact interface between zymogen ADAM17 and iRhom2. **Top:** Contact regions are highlighted with darker shades corresponding to their respective domains. **Bottom:** Conservation analysis is mapped onto the surface of the complex, with scores represented on a gradient from teal (variable) to maroon (conserved). **(B)** Zoomed-in view of the transmembrane domain of ADAM17 (color-coded by conservation) and TM1 of iRhom2 (gray) at Site 3. Annotated ADAM17 amino acids have side chains depicted as sticks. **(C)** Charged surface representation of the iRhom2 homology domain (HD), colored to indicate acidic residues (red) and basic residues (blue), in proximity to the ADAM17 cysteine-rich domain (green). Specific ADAM17 amino acids are labeled, with side chains shown as sticks (Site 2). **(D)** Western blot analysis of ConA-enriched lysates transfected with the ADAM17 mutations used in Figures 4B-C. The zymogen and mature forms of ADAM17 are marked with arrows.

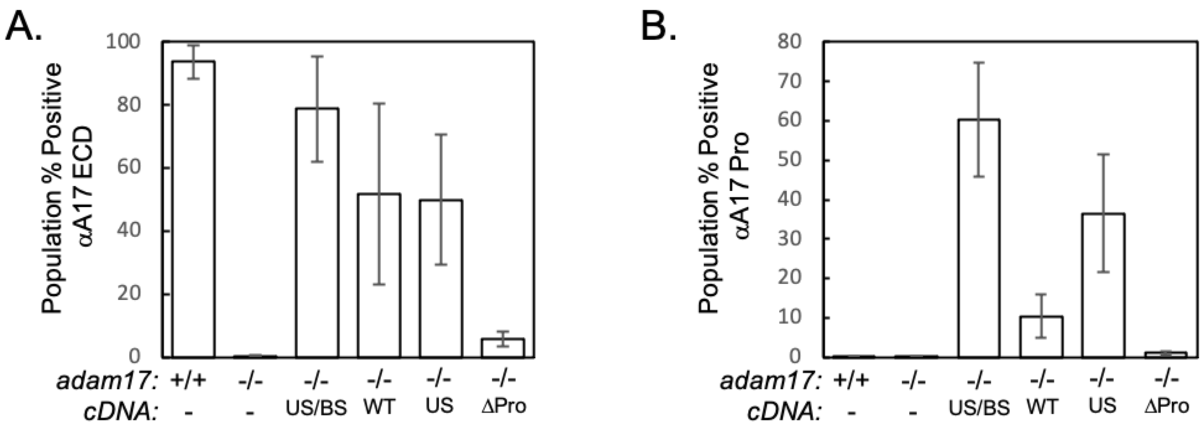


**Supplementary Figure 5: Quantification of ADAM17 and the Prodomain on the Cell Surface.** Quantification of flow cytometry analysis of the surface amount of (A) ADAM17 and (B) Prodomain on the cell surface of WT U2OS and adam17$^{-/-}$ cells transfected with ADAM17 expression constructs used in Figure 5. Error bars represent the mean ± SD from *N = 3* independent experiments.
